# Rapamycin reverts cavernoma endothelial cell phenotype and when combined with lapatinib ameliorates chronic lesions

**DOI:** 10.1101/2025.11.03.686432

**Authors:** Mar García-Colomer, José E Martínez, Luis Diaz-Gomez, Miriam Sartages, Eva M. Esquinas-Román, Cristina Riobello, David Martínez-Delgado, Diego González-Pérez, Aurora Gómez-Durán, Miguel Fidalgo, Marta Varela-Rey, Celia M Pombo, Juan Zalvide

**Affiliations:** Department of Physiology, Centro Singular de Medicina Molecular e Enfermedades Crónicas (CiMUS) and Instituto Sanitario de Santiago de Compostela (IDIS), Universidade de Santiago de Compostela, (USC), Santiago de Compostela, 15782, Spain; Department of Pharmacology, Pharmacy, and Pharmaceutical Technology, Instituto de Materiales (iMATUS), and IDIS, Universidade de Santiago de Compostela, Santiago de Compostela, 15782 Spain; Department of Biochemistry and Molecular Biology, CiMUS and IDIS, USC, Spain

## Abstract

This study investigates the impact of rapamycin and propranolol on cerebral cavernous malformations (CCMs). Employing an unbiased transcriptomic analysis, we aimed to comprehensively elucidate the molecular mechanisms underlying these drug effects in Mouse Brain Microvascular Endothelial Cells (mBMEC) deficient in *Ccm3*.

While propranolol shows limited efficacy in modulating the CCM transcriptomic phenotype in mBMEC, rapamycin demonstrates a higher impact. Rapamycin reverses gene expression changes induced by *Ccm3* deficiency, restoring *Klf2/4*-dependent genes like *Nos3*, *Adamts1*, and *Thbs1*. Notably, we observed a reduction in KLF2 protein levels in *Ccm3* KO cells treated with rapamycin.

Critically, *in vivo* experiments demonstrate that a combination of rapamycin and lapatinib effectively reduces lesion volume in a chronic CCM model. This finding is particularly noteworthy as it suggests a potential treatment strategy for existing lesions.

In summary, our work describes a new mechanism for the effects of rapamycin in *Ccm3*- deficient cells and identifies a new drug combination in the treatment of cavernomas.

## Introduction

Cerebral cavernous malformations (CCMs) are vascular lesions, predominantly occurring in the central nervous system and associated with recurrent hemorrhages and seizures (1, 2). CCMs are among the most common cerebral vascular malformations, especially in young people (it is estimated that 0.5% of the population has at least one CCM) (3, 4). Most CCMs are sporadic, but they can also have a genetic basis caused by inactivating mutations in one of the three CCM genes: *KRIT1/CCM1*, *OSM/CCM2* and *PDCD10/CCM3* (4). Patients with this familial cavernomatosis (fCCM) have multiple CCMs, which increases the severity of their condition. The products of these CCM genes form a ternary complex which binds to MEKK3, resulting in the inhibition of the MEKK3/MEK5/ERK5 axis and downregulation of the *KLF2* and *KLF4* transcription factors. Dysregulation of *KLF2* and *KLF4* in CCM mutant cells is essential for cavernoma development (5).

The current treatment for CCMs relies on surgical intervention for accessible lesions in the brain and spinal cord. However, for deep-seated CCM lesions and for fCCMs, when many lesions may appear, novel pharmacological interventions are clearly needed, and their development is a long-standing goal of translational CCM research. As a result of the studies on the function of CCM genes and cavernoma pathogenesis, several possible drug treatments have been proposed, among them statins, known to inhibit Rho activity, which is deregulated in CCM-deficient cells, especially in *Ccm1*- and *Ccm2*-deficient endothelium (6, 7); sulindac, which inhibits transcriptional β-catenin activity(8); the multikinase inhibitor sorafenib, which inhibits excessive angiogenesis (9); the free radical scavenger tempol (10); intestinal microbiome modification to alter Toll-mediated signaling (11); vitamin D and indirubin-3-monoxime, identified in a systematic pharmacological screen (12); lapatinib, because of the increase of EGFR in *Ccm3*- deficient endothelial cells (13); or rapamycin, based on the stimulated mTOR activity in CCM lesions (14). All these treatments diminish one or several of the effects of CCM deficiency *in vitro*, and some of them diminish the number of lesions that arise in animal models of cavernoma development, as long as they are given before their appearance or growth. Despite promising preclinical results, no clinically significant effect has been found in the two trials performed so far, a pilot study studying possible effects of by simvastatin on lesion permeability (15) and a blinded study on the effects of atorvastatin on lesion bleeding (16). Also, the phase 2 trial using the drug REC-99 (NCT05085561) has recently been discontinued because of lack of clinical benefits.

Interestingly, the largest number of clinical trials proposed for CCMs involve the non-selective β-blocker propranolol (trials NCT03589014, NCT03474614 and NCT03523650), which has been selected due to its known effect on infantile hemangiomas (17). Another treatment that has been proposed recently is rapamycin. This was initially based on the seminal observation that *Pik3ca* gain of function (GOF) mutations fueled growth of cavernoma lesions and was reinforced by the fact that CCM-deficient endothelial cells without *PIK3CA* mutations also had increased phosphorylation of the mTOR target S6 (14).

Propranolol treatment has been shown to reduce lesion burden in chronic murine models of cavernoma development and restore barrier function when initiated before or during lesion growth (18). Furthermore, while meta-analyses of clinical studies have not demonstrated a statistically significant protective effect of β-blockers (including propranolol) in preventing intracerebral hemorrhage or focal neurologic deficits in individuals with CCM, a phase II clinical trial suggested a potential benefit in preventing cavernoma development, although the study was not powered to achieve statistical significance (19). Despite these indications of activity, the clinical efficacy of propranolol remains unclear. Addressing this uncertainty requires further research, as the cellular mechanism of propranolol’s effect on CCM development is unknown, including the specific cell type in which it exerts its effects. Several studies have described the effects of propranolol on CCM-deficient endothelial cells, but at a concentration of 100 µM (20), which is significantly higher than the estimated *in vivo* concentration after propranolol treatment, even in the brain (no more than 10 µM). Moreover, treatment of mice with 15 mg/kg.day of propranolol did not affect lesion size or hemorrhage in already established cavernomas, at least under the treatment regimen used (20).

As stated above, the combination of the detection of *Pik3ca* GOF mutations in large cavernomas and enhanced mTOR activity in CCM-deficient cells makes mTOR-inhibiting drugs such as rapamycin good candidates for pharmacological treatment of cavernomas. Enhanced activity of mTOR has been shown in *CCM1*-deficient HuVECs (14) and in progenitor endothelial cells in *in vivo* lesions (21). Its inhibition with rapamycin reverts some of the effects of CCM loss in the latter, without affecting the mRNA overexpression of *Klf2* or *Klf4*, two genes that are important for cavernoma pathogenesis. It is unclear through what mechanism rapamycin affects the CCM phenotype, although it is believed that it acts specifically on progenitor endothelial cells (21).

In agreement with *in vitro* effects, treatment with rapamycin *in vivo* inhibits cavernoma development in several preclinical models (14, 21) However, the effects of rapamycin are not clear when a chronic model of CCM development and lower doses of the drug are used (22), suggesting that our understanding of the effects of rapamycin on CCM-deficient endothelial cells are not complete.

In this study, we investigated the effects of propranolol and rapamycin on *Ccm3*-deficient mBMEC. While neither treatment reversed known effects of CCM deficiency on protein membrane distribution, our unbiased transcriptomic analysis of drug effects revealed a key finding: rapamycin reverses the expression of genes dysregulated by *Ccm3* loss. In stark contrast, propranolol failed to demonstrate any significant effect in reversing the *Ccm3*-deficient phenotype, even under unbiased RNA-seq analysis. This observation suggests that if propranolol exhibits therapeutic effects in CCMs, it is unlikely to be through directly reversing the CCM phenotype in endothelial cells.

Importantly, we also present novel evidence that rapamycin, in combination with the tyrosine kinase inhibitor lapatinib, significantly reduces lesion volume in a chronic model of cavernomas. This finding underscores the potential of drug combinations as a promising therapeutic strategy for cavernomas, warranting further investigation.

## Results

To investigate the effects of drugs proposed for cavernoma treatment, we used mouse brain microvascular endothelial cells (mBMEC), a cellular model that closely mimics the capillary/venous endothelium from which these lesions originate. We employed mice carrying a floxed *Ccm3* allele (23) that express a tamoxifen-inducible cre recombinase (*cdh5*-creERT2) (24). To generate *Ccm3*-deficient mBMEC, we inactivated *Ccm3* in endothelial cells at passage 1 by treating the cultures with 4-OH-tamoxifen (4-OHT) for 4 days (25), which significantly reduced levels of *Ccm3* mRNA and protein compared to untreated cells (figure S1A and B). These *Ccm3*-deficient cells displayed characteristic effects of CCM deficiency, including elevated *Klf2* and *Klf4* mRNA levels (figure S1C) and reduced junctional levels of VE-cadherin and β-catenin (figure S1D). The presence of these established CCM-related phenotypes in our *Ccm3*-deficient mBMEC validates the suitability of this model for further investigating the effects of propranolol and rapamycin on CCM pathogenesis.

To further characterize the effects of *Ccm3* deficiency in mBMEC, we subjected cells to RNA-seq analysis. *Ccm3* inactivation deregulated 168 genes (figure S2A and table S1). Analysis of the differentially expressed genes (figure S2B) showed enrichment in biological processes that have been reported to be related to cavernomas, such as angiogenesis, cell adhesion, cell migration or extracellular matrix organization (26), which confirms that mBMEC are a good model to study the effects of CCM deficiency.

We sought to elucidate the specific effects of propranolol and rapamycin within this defined cellular model of CCM. We treated cells for 24 hours with 10 µM propranolol, based on its plasma levels in treated hypertensive patients and the distribution of the drug in brain tissue (27, 28). For rapamycin, we treated cells for 24 hours at a 100 nM concentration, based on the peak and trough levels attained in plasma after mice dosing (29–31). First, we analyzed the localization of VE-cadherin and β-catenin. VE-cadherin distribution was not affected by treatments in *Ccm3* KO cells, while a 24-hour exposure to rapamycin diminished its junctional localization in WT cells (figure 1A). β-catenin distribution to the cell junctions was also not affected by treatments in *Ccm3* KO cells, but, surprisingly, both drugs reduced its amount in wild-type cells after a 24-hour exposure, making them more similar to *Ccm3*-deficient cells at least in this regard (figure 1B). We concluded that neither rapamycin nor propranolol recovered the abnormal distribution of VE cadherin or β -catenin in *Ccm3*-deficient mBMEC.

**Figure 1.**
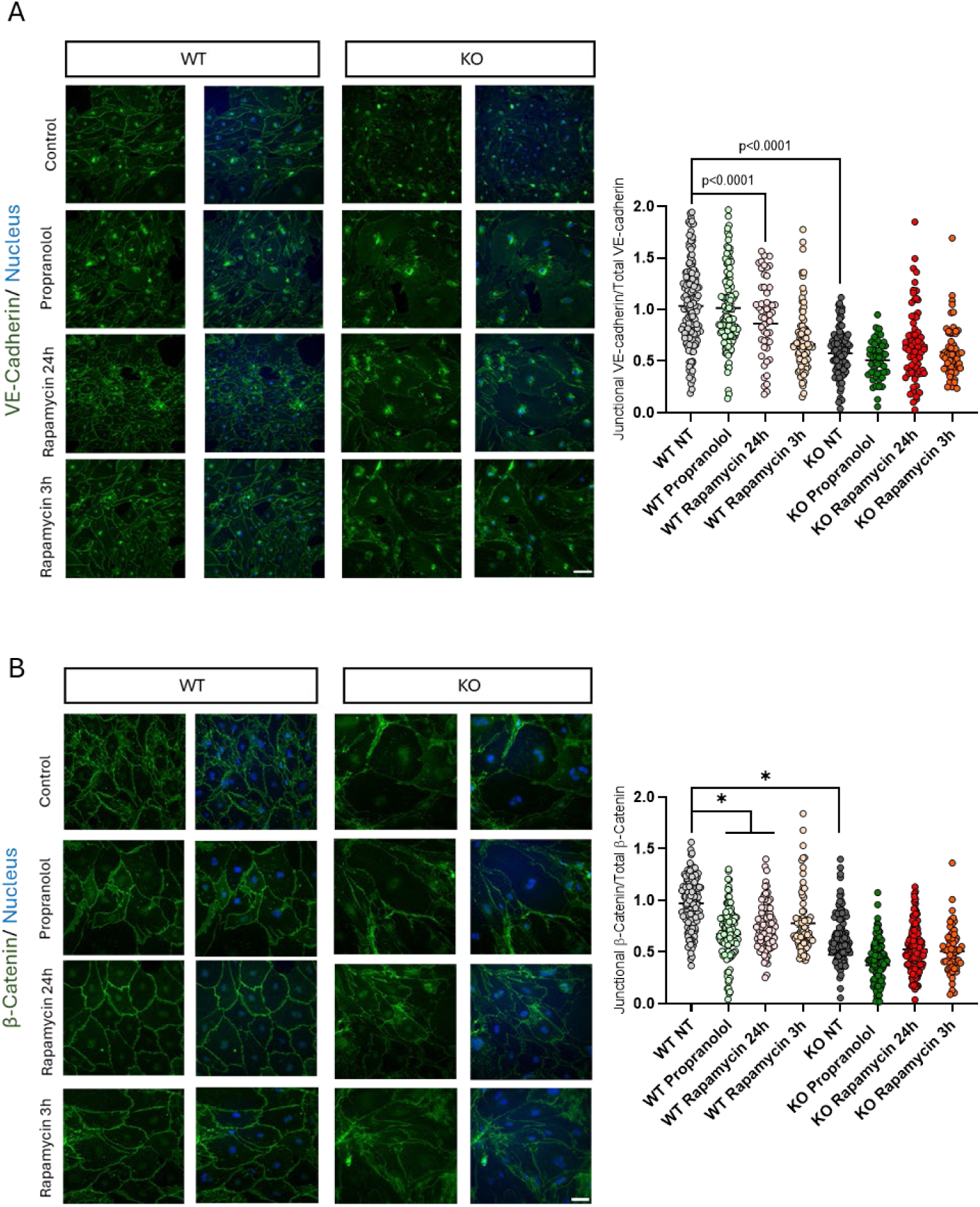
Rapamycin or propranolol do not recover abnormal distribution of VE-cadherin or β-catenin in *Ccm3* KO mBMEC. **A**. Distribution of VE-cadherin. WT or KO mBMEC were either left untreated or treated with 10 µM propranolol for 24 h, 100 nM rapamycin for 24 h, or 100 nM rapamycin for 3 h, and an immunofluorescence performed for VE-cadherin, counterstaining with DAPI. Upper panels: representative photographs of the immunofluorescence with staining for VE-cadherin (green) and DNA (blue). Scale bar is 50 µm. Graph shows the ratio of junctional/total VE-cadherin after each treatment, referenced to untreated WT mBMEC, of >100 cells per treatment from 4 independent biological replicates (cell cultures). Each cell culture was obtained from a pool of 3 to 4 brains. P-values are from ANOVA analysis with a Tukey’s multiple comparison test. **B**. Distribution of β-catenin in WT and KO mBMEC. Upper panels: representative photographs of the immunofluorescence with staining for β-catenin (green) and DNA (blue). Scale bar is 50 µm. Graph shows the ratio of junctional/total β-catenin after each treatment, referenced to untreated WT mBMEC, of >100 cells per treatment from 4 independent biological replicates (cell cultures). Each cell culture was obtained from a pool of 3 to 4 brains. P-values are from ANOVA analysis with a Tukey’s multiple comparison test.

To study the effect of both treatments in an unbiased manner, we performed an RNA-seq analysis of *Ccm3* KO cells after treatment and analyzed their effect on the 168 genes that were differentially expressed in KO vs WT cells, which reflected the effect of *Ccm3* loss on brain endothelial cells. Treatment with propranolol affected the expression of several of these genes (figure 2A). However, when a principal component analysis (PCA) was performed comparing WT, untreated and treated KO cells, the effect of propranolol was shown not to revert the effects of *Ccm3* deficiency but acted along a different component (figure 2B). In contrast, rapamycin partially recovered the displacement induced by the loss of *Ccm3*, suggesting a normalization of gene expression and thus a possible therapeutic effect in these cells (figure 2D). This is also reflected in the heatmap of the DEGs, where KO cells treated with rapamycin cluster apart from untreated KO cells (figure 2C).

**Figure 2.**
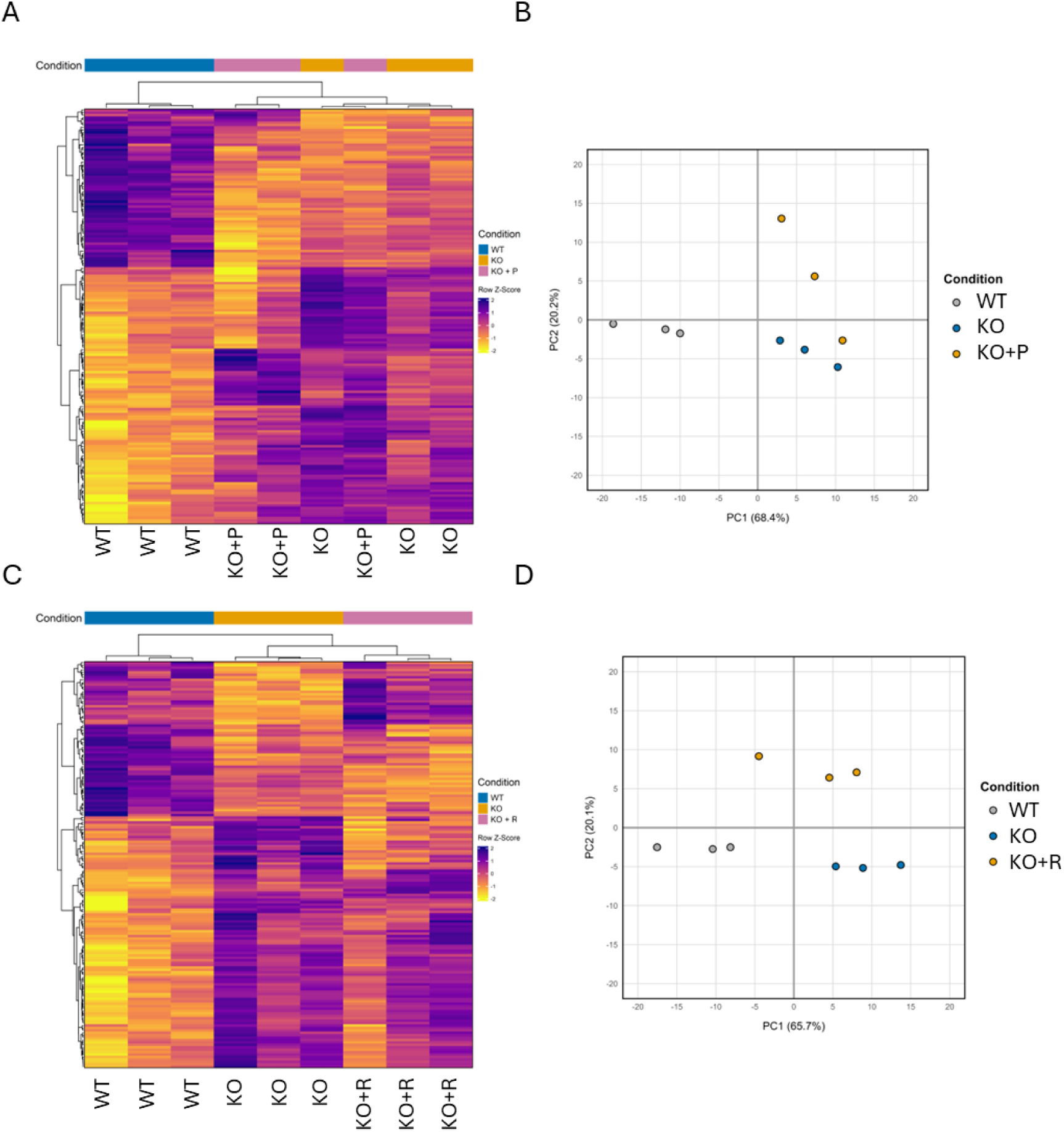
Rapamycin, but not propranolol, can partially revert the transcriptional effect of *Ccm3* inactivation in mBMEC. WT and KO mBMEC were left untreated or treated with 10 µM propranolol for 24 h (KO+P) or 100 nM rapamycin for 24 h (KO+R). RNAseq analysis was performed, and their expression profile was compared using genes differentially expressed in KO vs WT mBMEC. Three independent biological replicates (cell cultures) were analyzed in all cases. Each cell cultures was obtained from a pool of 3 to 4 brains. **A.** Heatmap analysis of WT, KO, and KO+P, showing that KO+ P cells cluster with KO cells. **B.** Principal component analysis of WT, KO, and KO+P mBMEC, showing a displacement of KO vs WT mBMEC in the horizontal axis that is not reverted by treatment with propranolol. **C.** Heatmap analysis of WT, KO, and KO+R, showing that treated KO+R mBMEC no longer cluster with KO mBMEC. **D.** Principal component analysis of WT, KO, and KO+R mBMEC, showing a displacement of KO WT mBMEC in the horizontal axis that is partially reverted by treatment with rapamycin.

In-depth analysis of the genes recovered by rapamycin showed that they included some of the genes known to be dependent on *Klf2/4* transcription factors, which are deregulated in CCM deficient cells. This included genes such as *Adamts1*, overexpressed in CCM-deficient cells (32, 33), or *Thbs1*, downregulated in CCM-deficient deficient cells (26). To confirm this effect, we performed RT-qPCRs in cell preparations different from those used for RNA-seq. *Adamts1* was confirmed as overexpressed and inhibited by rapamycin in *Ccm3* KO mBMEC (figure 3A), while *Thbs1* was downregulated in KO cells, with rapamycin recovering it (figure 3B). *Nos3*, a gene regulated by *Klf2/4* and important in CCM pathogenesis (34), was also overexpressed in KO cells and inhibited by rapamycin when checked by RT-qPCR (figure 3C). On the contrary, *Ccm3*, *Klf2* and *Klf4* were not affected by rapamycin treatment in *Ccm3* KO cells (figures 3D, E and F), which is consistent with its described effects in other CCM-deficient endothelial cells (14). Contrary to the effects of rapamycin, and in accordance to the RNA-seq results, propranolol did not influence the expression of any of these genes.

**Figure 3.**
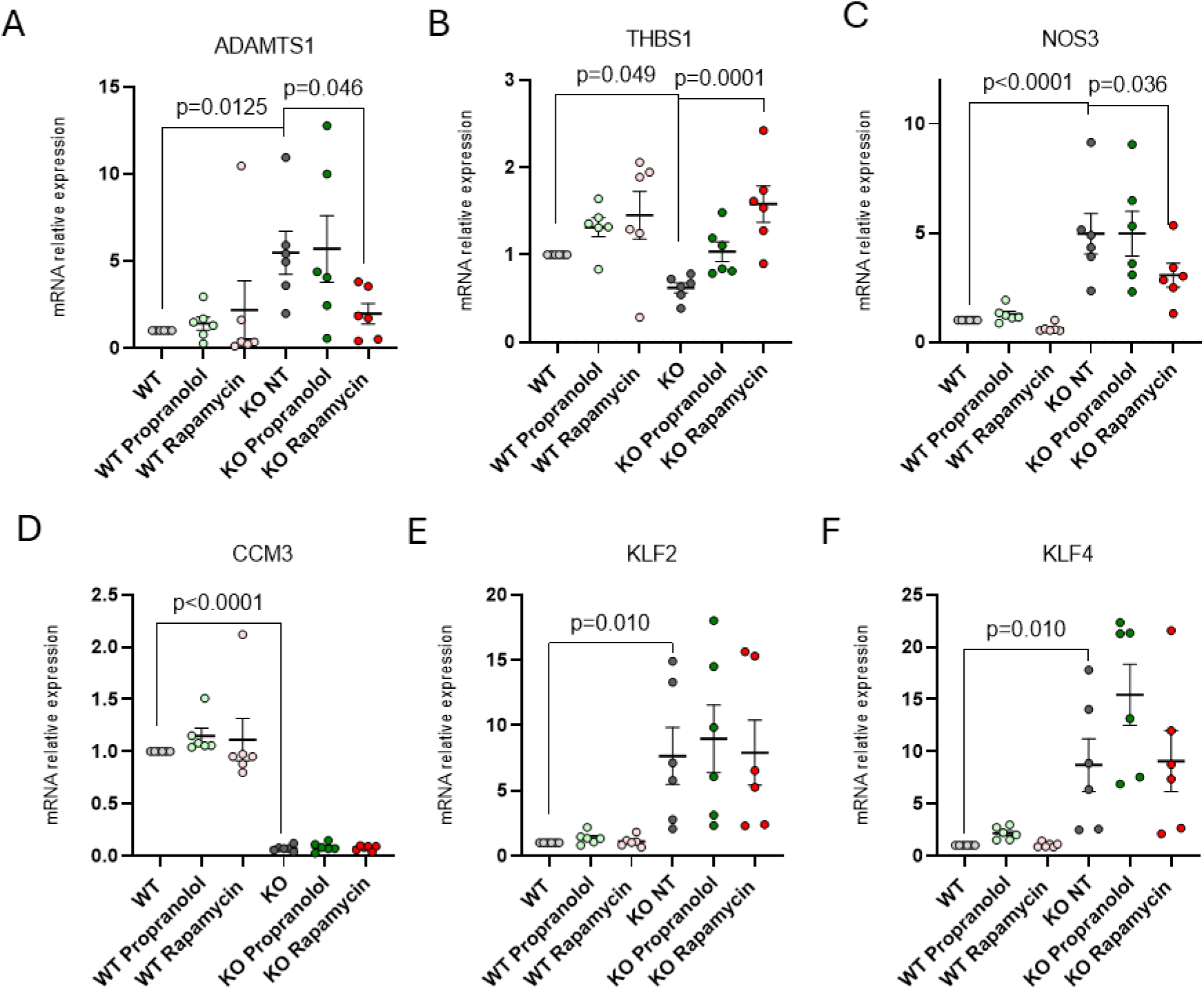
Rapamycin, but not propranolol, reverts the expression of genes dependent on *Klf2/4*, without affecting *Klf2* or *Klf4* mRNA expression. WT or KO mBMEC were either left untreated or treated with 10 µM propranolol for 24 h or 100 nM rapamycin for 24 h. RNA was extracted and RT-qPCR was performed using GAPDH as a reference gene. Each graph shows the average and SEM of mRNA relative expression and the individual values of six independent biological replicates (cell cultures). Each cell culture was obtained from a pool of 3 to 4 brains. P-values are from ANOVA analysis with a Tukey’s multiple comparison test. **A.** Expression of *Nos3*, **B.** Expression of *Adamts1*, **C.** Expression of *Thbs1*, **D.** Expression of *Ccm3*, **E.** Expression of *Klf2*, **F.** Expression of *Klf4*.

Overexpression of *Nos3* is one of the hallmarks of cavernous malformations, and it is important for lesion development through a feedback loop involving NO production by cavernoma endothelial cells and the ensuing secretion of VEGF by astrocytes that contributes to lesion enlargement (34). Thus, we wanted to see if the effect of rapamycin on *Nos3* mRNA was reflected at the protein level. As seen in figure 4A and B, both eNOS protein levels, and its phosphorylation in serine 1177 were stimulated in *Ccm3*-deficient cells, and this was reverted when they were treated with rapamycin for 24 hours. The same western blot shows that *Ccm3*-deficient cells had a tendency to have higher levels of S6 phosphorylation with no change in Akt phosphorylation in S473, which, together with a higher S6K1 phosphorylation in Serine 371 (figure 4D), is in agreement with the effects of *Ccm1* KO in brain endothelial cells and HuVECs, as reported by Ren et al (14). Rapamycin acted as expected both in WT and KO cells, inhibiting S6 phosphorylation after 3 and 24 hours of treatment, and diminishing AKT phosphorylation in S473 only after 24 hours.

**Figure 4.**
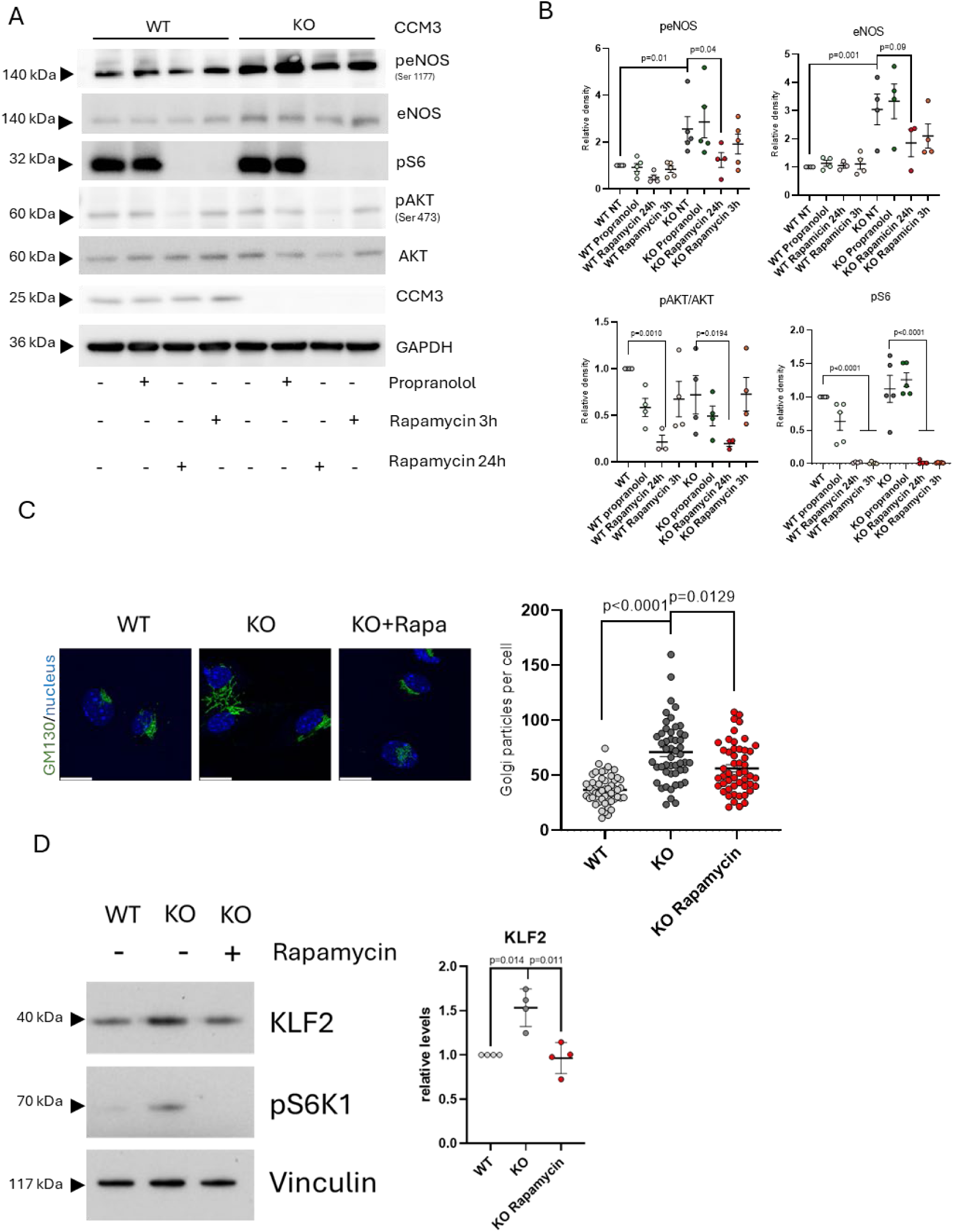
Effect of rapamycin on eNOS, Golgi dispersion and KLF2 protein levels. **A.** Western blot of mBMEC proteins after treatments. Extracts were prepared from WT and KO mBMEC treated with 100 nM rapamycin for 3 or 24 hours, or with 10 µM propranolol for 24 hours, and western blots performed for total eNOS, eNOS phosphorylated in Ser 1177 (peNOS), phosphorylated S6 (pS6), AKT phosphorylated in Ser 473 (pAKT), total AKT, *Ccm3*, with GAPDH as a loading control. **B.** Average and SEM of the intensity of peNOS, eNOS, pAKT/AKT ratio and pS6 and from four independent biological replicates (cell cultures). Each cell culture was obtained from a pool of 3 to 4 brains. P- values are from ANOVA analysis with a Tukey’s multiple comparison test. **C.** Rapamycin treatment reverts Golgi dispersion in *Ccm3*-deficient mBMEC. WT and KO mBMEC were untreated or treated with 100 nM rapamycin for 24 hours and stained for GM130 to visualize Golgi. Left panels: Photographs of untreated WT cells, and KO cells untreated or treated with rapamycin. Scale bar is 20 µm. Right graph: Golgi fragments per cell of >100 cells per treatment from 3 independent biological replicates (cell cultures). Each cell culture was obtained from a pool of 3 to 4 brains. P-values are from ANOVA analysis with a Tukey’s multiple comparison test. **D.** Rapamycin diminishes KLF2 protein levels in *Ccm3* deficient cells. KO and WT mBMEC were left untreated or treated with 100 nM rapamycin for 24 hours. Western blots were performed for KLF2, vinculin and pS6K1. Left panel, representative western blot. Right graph, average and SEM of corrected intensity of KLF2 of western blots from four biological replicates (cell cultures). Each cell culture was obtained from a pool of 3 to 4 brains. P-values are from ANOVA analysis with a Tukey’s multiple comparison test.

Another important functional consequence of *Ccm3* deficiency in endothelial cells is the abnormal dispersion of the Golgi apparatus, which is linked to their impaired ability to polarize during cell migration (35). Notably, rapamycin partially reversed this dispersion in *Ccm3* KO mBMEC (figure 4C).

The effect of rapamycin on known transcriptional targets of *Klf2* and *Klf4*, together with its lack of effect on the expression of these transcription factors at the mRNA level, hints at a possible post-transcriptional effect of the drug on *Klf2* and/or *Klf4*, probably through the known effects of rapamycin on translation. However, there is no apparent effect on *Klf4*, as its protein levels have been shown not to change with rapamycin in *Ccm1*- deficient endothelial cells, both *in vivo* and *in vitro* (14). Intriguingly, in a different setting, Wang et al have shown that rapamycin inhibits *Klf2* by post-transcriptional mechanisms in endothelial cells (36). When we checked KLF2 protein levels in *Ccm3* KO cells, we saw that rapamycin treatment could diminish KLF2 protein (figure 4D), even if it did not inhibit its mRNA. This shows that rapamycin inhibits KLF2 even without affecting its mRNA levels, and points to a direct mechanism for the effect of the drug on KLF2/4 targets in *Ccm3* KO cells.

The effects of rapamycin in the reversal of alterations induced by the lack of *Ccm3* reinforce the idea that it could be used for treatment both of growing and mature cavernoma lesions, either alone or in combination with other agents. Rapamycin has been shown to prevent the development of cavernomas using animal models of the disease, both in lesions with only mutations in a CCM gene and in those harboring also *Pik3ca* mutations (14). However, concerns have been raised on these results based on the high doses of rapamycin used, which may have achieved supratherapeutic levels that cannot be safely tolerated in humans (31). The trough levels of rapamycin that are usually targeted in plasma for benign conditions are 5-15 ng/mL (31, 37). Thus, we decided to induce cavernoma in neonatal *Cdh5* (PAC)-CreERT2/*Pdcd10*^fl/fl^ mice (24) and administer 1.5 mg/Kg rapamycin intraperitoneally, a lower dose than the 2 mg/Kg used by Bishu *et al* to attain trough levels of 13.8 ng/mL (37) (figure 5A). Under this regime, rapamycin could inhibit the development of cavernoma lesions, although it did not completely prevent them, as happens when higher doses are used (14, 21, 38) (figure 5B and 5C).

**Figure 5.**
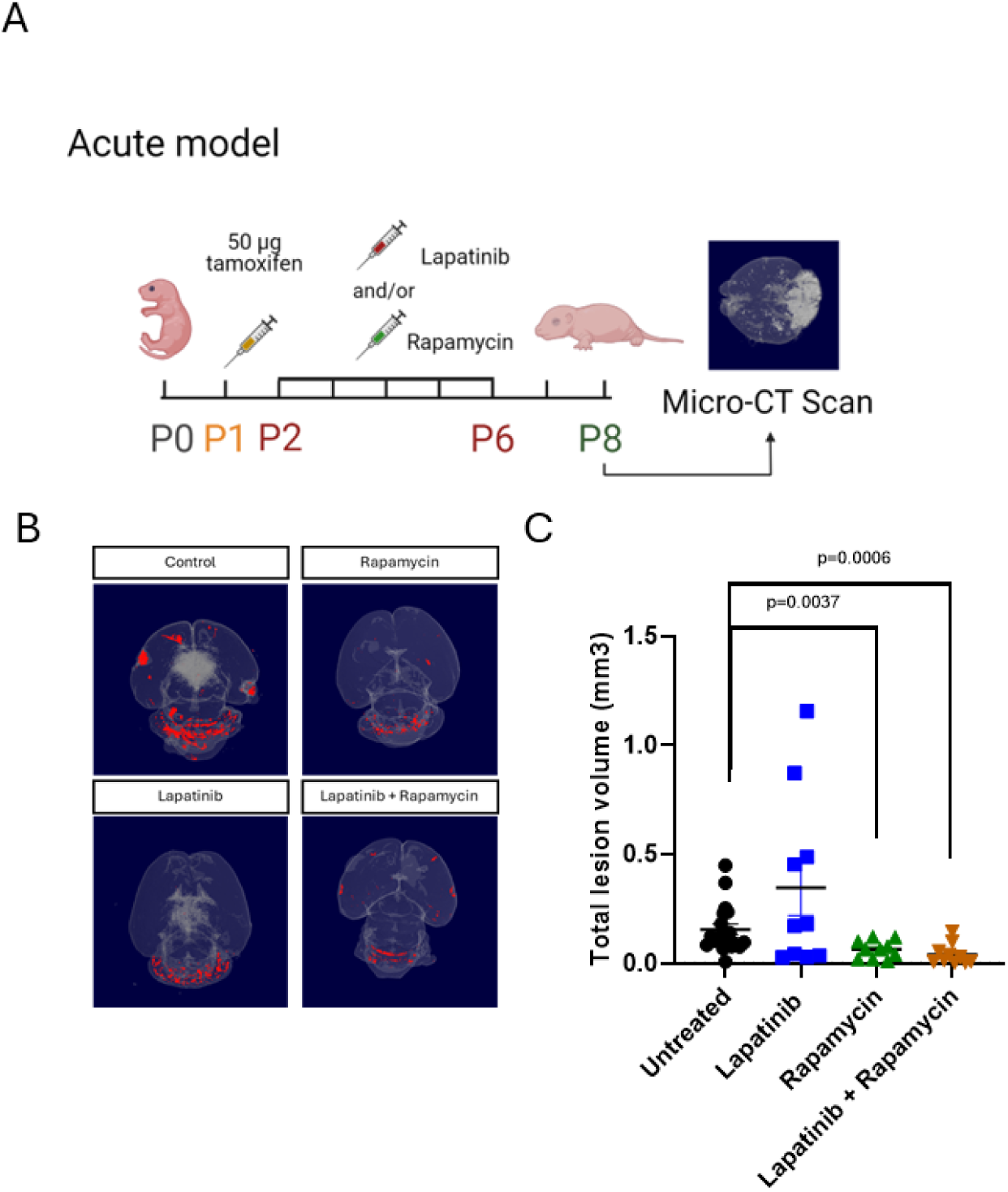
Rapamycin, but not Lapatinib, can inhibit acute cavernoma development. **A.** Scheme showing the induction and treatment of cavernomas. 50 µg tamoxifen were administered to newborn Cdh5 (PAC)-CreERT2/PDCD10^fl/fl^ mice at P1. Rapamycin and/or Lapatinib were injected at days P2 to P6, and mice were euthanized at P8, brains extracted and analyzed by micro-CT. **B.** Micro-CT images of brains of P8 mice with induced cavernomas untreated, or treated with Lapatinib, Rapamycin, or Lapatinib + Rapamycin. **C.** Volume of cavernomas untreated mice (n=19 mice), or mice treated with Lapatinib (n=10 mice), Rapamycin (n=10 mice), or Lapatinib + Rapamycin (n=10 mice). Total volumes of cavernomas per brain were calculated from micro-CT images. P-values are from Welch’s ANOVA test followed by a Dunnett’s T3 multiple comparisons test. Panel A was created with biorender.com.

We then considered the possibility of using rapamycin in combination with other agents for the treatment of cavernomas. Since propranolol had no apparent effect in our cellular system, we turned to the HER2/EGFR inhibitor lapatinib, as we had previously shown that *Ccm3*-deficient endothelial cells have higher levels of EGF receptor, and are more susceptible to apoptosis after its inhibition with lapatinib *in vitro* (13). Further, it is known that high levels of PI3K activity in tumors make them resistant to EGF receptor inhibition and rapamycin enhances the effect of EGFR inhibition in breast cancer cells (39). Treatment with 300 mg/Kg lapatinib alone did not significantly prevent cavernoma development. However, the combined rapamycin/lapatinib treatment resulted in a trend towards lower volume of the cerebellum occupied with lesions as compared with the already low volume of cavernomas in mice treated with rapamycin alone (figure 5B and 5C). This hinted at the possibility that it could be a good strategy to inhibit the EGF receptor family in combination with rapamycin treatment for the handling of cavernomatosis, and that it could have a greater effect in situations where rapamycin alone is less effective, such as cavernomas that have already developed.

Being able to inhibit lesions that have already developed would be an optimal strategy for treatment of cavernomas. Unfortunately, treatment of chronic lesions with pharmacologically relevant doses of rapamycin does not result in lesion improvement (22). We wanted to see if the addition of lapatinib could help reduce lesion volume of more mature lesions. We used a model of chronic cavernoma developed for the same animals used for acute cavernoma development by Malinverno *et al* (40), and treated them with an injection of rapamycin alone or in combination with lapatinib every 24 hours from postnatal day 30 (P30) to P39, measuring cavernoma volume at P40 (figure 6A). Consistent with recent reports (22), rapamycin alone did not affect the evolution of lesions in this model but, surprisingly, combined treatment of rapamycin and lapatinib had an effect on the volume occupied by lesions in these animals (figure 6B), suggesting that the combination of rapamycin with inhibition of tyrosine kinase receptors may be especially useful to treat established cavernomas.

**Figure 6.**
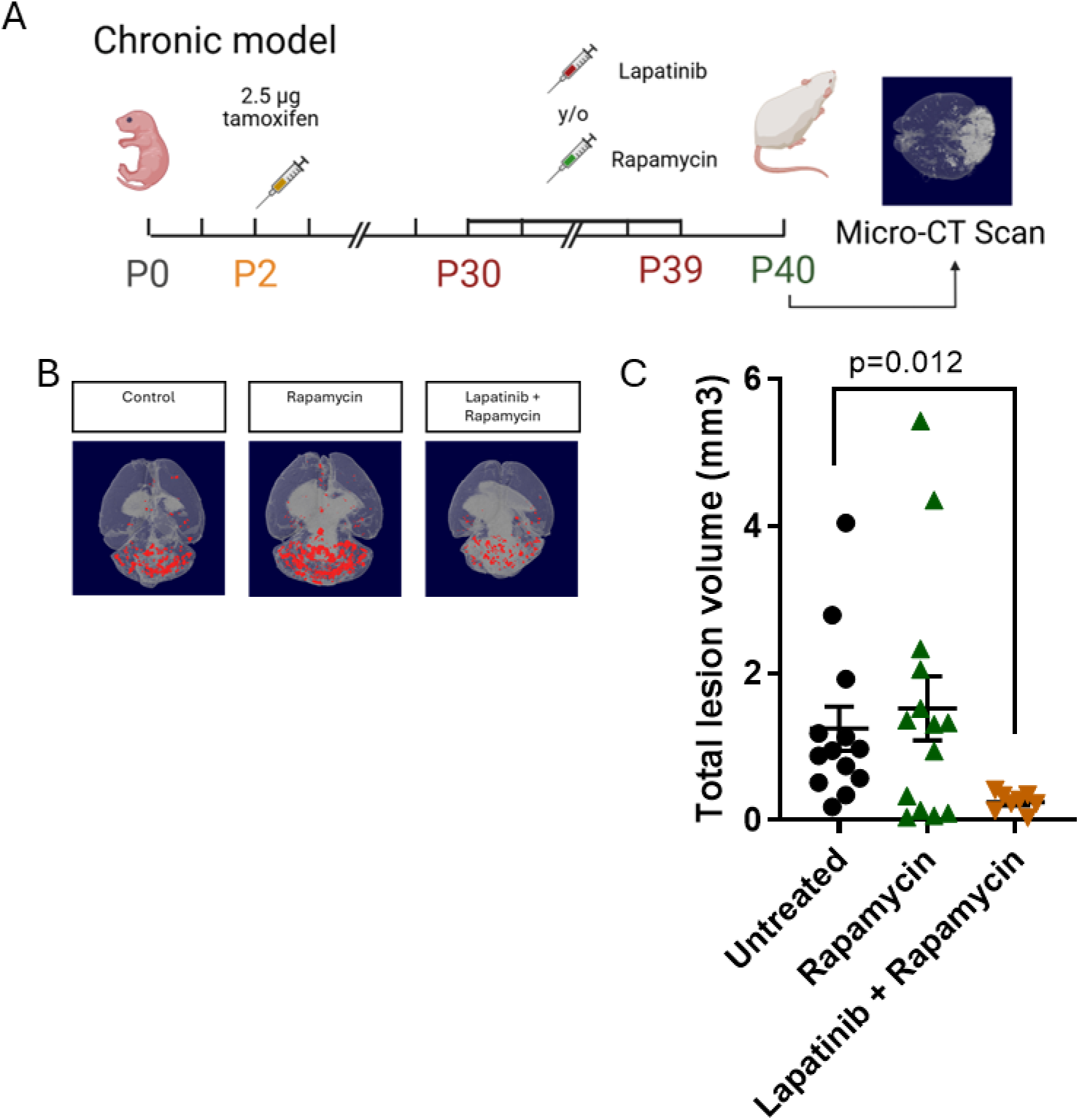
Rapamycin/Lapatinib combination can diminish cavernoma volume when used after cavernoma development. **A.** Scheme showing the induction and treatment of cavernomas. 2.5 µg tamoxifen were administered to Cdh5 (PAC)-CreERT2/PDCD10^fl/fl^ mice at P2. Rapamycin or Rapamycin + Lapatinib were injected at days P30 to P39, and mice were euthanized at P40, brains extracted and analyzed by micro-CT. **B.** Micro-CT images of brains of P30 mice with induced cavernomas untreated, or treated with Rapamycin or Lapatinib + Rapamycin. **C.** Volume of cavernomas in untreated mice (n= 13 mice), or mice treated with Rapamycin (n=14 mice) or Lapatinib + Rapamycin (n=8 mice). Total volumes of cavernomas per brain were calculated from micro-CT images. P-values are from Welch’s ANOVA test followed by a Dunnett’s T3 multiple comparisons test. Panel A was created with biorender.com.

## Discussion

We have investigated the effects of two proposed cavernoma treatments, rapamycin and propranolol, on the biology of *Ccm3*-deficient brain endothelial cells. To ensure an unbiased approach, we analyzed the impact of both drugs on the transcriptomic signature of *Ccm3* KO brain endothelial cells, assessing their ability to revert it toward that of WT cells. Using this strategy, we discovered that propranolol exhibits no discernible effect on the transcriptional phenotype induced by *Ccm3* loss in our model with the doses and treatment time used. In contrast, rapamycin significantly reverses the transcriptomic signature triggered by *Ccm3* deficiency.

Rapamycin was first proposed as a possible treatment for cavernomas by Ren et al (14), based on their groundbreaking discovery of activating *Pik3ca* mutations in cavernomas and the observation that *Ccm1* loss or *KLF4* overexpression increased mTORC1 activity in cavernoma lesions and HuVECs, an effect which has been described as mediated by caveolin 1(21). Our cellular model effectively recapitulates the effects of CCM deficiency on mTOR activity in endothelial cells. Importantly, despite the absence of *Pik3ca* activating mutations, we observed signs of enhanced mTOR activity in *Ccm3*-deficient mBMEC compared to *Ccm3* WT cells, as evidenced by S6K1 phosphorylation and a trend towards S6 phosphorylation, while AKT phosphorylation remained unaffected. These effects mirror those resulting from *Ccm1* loss (14).

Using this model, we show that rapamycin significantly impacts the expression of genes known to be upregulated in CCM-deficient endothelial cells, such as *Nos3*, *Adamts1* and *Thbs1*. Moreover, the deregulated expression of these genes in KO cells is known to be dependent on the transcription factors *Klf2* and *Klf4*. Importantly, we also show that rapamycin, despite not affecting its mRNA levels, diminishes KLF2 protein. This suggests a mechanism for the restoration of expression of *Klf2/4* transcriptional targets in *Ccm3* KO cells. This specific effect on KLF2 is particularly noteworthy, as rapamycin is known not to affect KLF4 protein levels (14). Interestingly, while rapamycin treatment influences several pathways altered in cavernomas, it does not affect them all. Specifically, rapamycin does not revert the altered distribution of β-catenin or VE cadherin in the membrane of *Ccm3*-deficient cells, suggesting these alterations are not dependent on KLF2 overexpression.

The transcriptional effects of rapamycin have important functional consequences. *Nos3* overexpression and the ensuing overproduction of NO in CCM-deficient endothelial cells is known to be important in cavernoma pathogenesis (34), whereas inhibition of *Thbs1* also favors cavernoma development (26). We show that rapamycin can diminish the amount of the *Nos3* gene product, eNOS and its phosphorylation, and can revert Golgi dispersion in CCM-deficient cells. This underscores the potential of rapamycin as a component of a pharmacological strategy to treat cavernomas, even in the absence of *PIK3CA* mutations.

Importantly, in this study we provide compelling evidence that rapamycin, when combined with a drug possessing a distinct mechanism of action such as lapatinib, can effectively improve chronic cavernoma lesions. With the cavernoma model developed by Malinverno et al. (40), we can begin treatment at postnatal day 30, 28 days after lesion induction, when cavernomas are known to be already detectable. This suggests that the combination of rapamycin and lapatinib, but not rapamycin alone, may improve cavernomas after they have already formed. A common characteristic of all proposed cavernoma treatments that have failed in clinical trials to date is their efficacy in preventing cavernoma development, rather than in improving existing cavernomas. However, in clinical settings, cavernomas are typically already detectable upon diagnosis. The activity of the rapamycin/lapatinib combination in improving cavernomas after 30 days of development makes this strategy worthy of further pursuit, including refinement in preclinical models and incorporating MRI exams in animals before and after treatment.

In contrast to rapamycin, propranolol exhibits no activity in reverting the phenotype of *Ccm3*-deficient cells in our model, both when analyzed in a directed manner (e.g., β- catenin or VE-cadherin distribution) and in an unbiased manner (RNA-seq analysis). These results diverge from reports demonstrating phenotypic effects of propranolol in CCM-deficient endothelial cells (18, 20, 41). However, in those experiments, the concentration of propranolol used was significantly higher (50 to 100 µM) and not readily achievable in clinical settings. One possibility is that propranolol has beneficial effects on CCM-deficient endothelium that do not involve reversing the CCM phenotype. Alternatively, the reported effects of propranolol on intracranial hemorrhage or focal neurological deficits in patients and animal models may be mediated by its actions in non-endothelial cells or on a specific subset of endothelial cells within a cavernoma that are not reflected in the mBMEC used in this study.

In summary, our study demonstrates that rapamycin effectively reverses key aspects of CCM3 deficiency in brain endothelial cells and, when combined with lapatinib, significantly reduces lesion volume in a chronic model. This highlights the potential of rapamycin-based combination therapies as a promising strategy for treating established CCM lesions, addressing a critical unmet need in cavernoma therapy.

## Methods

### Sex as a biological variable

Male and female mice were used in the experiments, as no sex differences have been reported in the development of cavernomas.

### Mouse experiments

All experiments involving animals were performed following the ARRIVE guidelines (42). Number of mice used in each experimental group are outlined in figure legends.

All mice used in this study, both for the isolation of brain endothelial cells and for *in vivo* treatments, were maintained in a C57BL/6J background from those developed by Elisabetta Dejana laboratory crossing *Pdcd10*^fl/fl^ mice (8) with *cdh5*(PAC)-CreERT2 mice (24), as described in (8).

#### Cavernoma induction

For acute cavernoma induction (acute model), pups were injected at postnatal day 1 (P1) intragastrically with 50 µg tamoxifen (Merck Life Sciences #T5648) to obtain *Ccm3*^iEC/iEC^ mice and induce CCM lesions.

To induce the chronic cavernoma lesions (chronic model), pups were injected at P2 intragastrically with 2.5 µg tamoxifen to obtain *Ccm3*^iEC/iEC^ mice.

#### Drug administration

Treatment for each individual animal within a litter was randomly assigned at birth independently of sex. Sample size was based on previously published reports on cavernoma treatments using the same animal model.

Rapamycin (MedChemExpress #HY-102129) was dissolved in DMSO and injected intraperitoneally at a dose of 1.5 mg/Kg. Lapatinib (MedChemExpress #HY-50898) was dissolved in DMSO and then diluted in PBS, either alone or in combination with rapamycin, and injected intraperitoneally at a dose of 300 mg/kg. Control animals were administered a volume of DMSO equal to that received by treated mice.

In the acute model, animals were injected intraperitoneally with rapamycin at postnatal day 2 (P2), lapatinib from P2 to P6, a combination of both drugs or vehicle from P2 to P6. Mice were euthanized at P8 by CO_2_ inhalation followed by cervical dislocation. For experiments with the chronic model, adult mice were injected intraperitoneally with rapamycin, lapatinib, rapamycin + lapatinib, or vehicle from P30 to P39, following the same dosage regime. Mice were euthanized at P40 by CO_2_ inhalation followed by cervical dislocation.

For analysis of cavernomas or isolation of mBMEC, animals were euthanized by CO_2_ inhalation followed by cervical dislocation.

#### Assessment of lesions

After euthanasia by CO_2_ inhalation followed by cervical dislocation brains were collected and fixed with formalin until staining with Lugol’s iodine (Merck Life Sci, #L6146) at 50% in water solution for 96 h (acute model) or 5 days (chronic model) under gentle agitation (43). Following incubation, micro-CT imaging was conducted using a Skyscan 1272 system (Bruker, Kontich, Belgium) at 100 kV and 100 μA, with a 10 μm pixel size, 1150 ms exposure time, and a 0.4° rotation step, using an Al 0.5 + Cu 0.038 filter. Image projections were reconstructed with NRecon software (Bruker) and rendered using CTVox (Bruker). Volumetric segmentation and quantification of total volume of CCM lesions in each animal were carried out using CTAn software (Bruker). All imaging and volume quantification were performed in a blinded manner, with reconstruction parameters kept constant throughout the analysis. Those litters in which the average of the cavernoma volume of all animals was less than 0.05 mm^3^ in acute induction or 0.5 mm^3^ in chronic induction were excluded. The experimental unit for analysis was the brain of non-excluded individual animals.

### Isolation of mBMEC and treatments

mBMEC were isolated from adult 8–10-week *Cdh5* (PAC)-CreERT2/*Pdcd10*^fl/fl^ mice. Mice were euthanized and brain tissue harvest and tissue homogenization and digestion performed as described by Czupalla (44). Then cells were resuspended and cultured as described by Assmann with slight modifications (45). After cell isolation, cells were seeded onto plates coated with collagen I (Merck Life Sciences #C3867) in DMEM-high glucose supplemented 15% v/v fetal bovine serum (Gibco™ #10270-106), 1% v/v Penicillin/Streptomycin/L-Glutamine (Corning 30-009-CI), 100 μg/mL heparin (Merck Life Sci #H3149) and 50 μg/ml endothelial cell growth supplement (ECGS; Merck Life Sci #E2759). Cells were maintained at 37°C in 95% air and 5% CO_2_.

After 24 hours, endothelial cells were selected using 4 μg/ml puromycin for 3 days, after which Cre recombinase activity was induced using 5 μg/ml 4-hydroxy-tamoxifen (Merck Life Sciences #SMC1666) over ten days.

mBMEC were treated, when they reached confluence, with either propranolol (Merck Life Sci #P0884) or rapamycin (#HY-102129) at concentrations and time points indicated in the corresponding figure legends.

### Immunofluorescence and quantification

mBMEC were grown on coverslips pretreated with collagen I, allowed to reach confluency and treated for the times indicated in the corresponding figure legends. Then, coverslips were fixed using 4% paraformaldehyde in PBS pH 7.4 at room temperature for 15 minutes and permeabilized with Triton X-100 0.5% in PBS. Cells were then processed for immunofluorescence using 10% FBS as blocking solution for 60 min.

The primary antibodies used for immunostaining are listed in table S2. Fluorescence-conjugated Alexa Fluor 488 secondary antibodies were used according to the primary antibody species (1:250, Invitrogen #A11001 or #A11008) and counterstained with DAPI (1:500). The coverslips were mounted with Fluoroshield (Merck Life Sci #F6182).

Images were acquired using Leica DM4-B and Leica K5 camera microscope or a Leica confocal microscope equipped with an HCX PL APO CS 63x/1.32 objective. Images were processed with the LAS X (Leica) and quantified using the ImageJ (National Institutes of Health) software. The junctional/total ratio of VE-cadherin and β-catenin was calculated according to Colás-Algora et al. (35). Golgi dispersion was quantified counting the number of Golgi fragments in a cell.

### Protein immunoblotting

Western blotting was performed by standard procedures after preparation of mBMEC extracts in cold radioimmunoprecipitation assay (RIPA) lysis buffer plus protease and phosphatase inhibitors. For KLF2 detection, lysis buffer was 80 mM Tris-HCl pH7.8; 10% glycerol; 2% SDS, 1% β-mercaptoethanol, plus protease and phosphatase inhibitors. Blots were developed using ECL® Chemiluminescent detection reagents (Thermo Fisher Scientific Inc., Waltham, MA, USA #32106) and ChemiDocTM MP (Biograd) or developed using Super RX-N Fuji medical X-Ray Films (Fujifilm #47410-19289). The software ImageJ (version 1.52s, National Institute of Health, Bethesda, MD, USA) was used to quantify the Western blot signals.

The primary antibodies used for western blot are listed in table S2. The following secondary HRP-conjugated antibodies were used: IgG mouse (H+L) 1 mg/ml (Invitrogen, #A16072) and IgG rabbit (H+L) 1 mg/ml (Invitrogen, #A16035).

### RNA extraction, sequencing and data analysis

Total RNAs were extracted using the Trizol reagent (life Technologies), according to the manufacturer’s protocol. For RNA-seq, RNA integrity and quantity of the samples was analyzed using a NanoDrop spectrophotometer followed by agarose gel validation.

For RT-qPCR RNA was reverse-transcribed using random hexamers, and the mRNA expression level was determined using PowerUp™ SYBR™ Green Master Mix (Thermo Fisher Scientific Inc., Waltham, MA, USA #A25778) using a StepOnePlus™ Real-Time PCR System. *Gapdh* mRNA levels were used as internal control, and the 2^-ΔΔCT^ method was used for analysis of the data. Each control value (*Ccm3* WT) was normalized to 1, and *Ccm3* KO values were relative to control.

Oligonucleotides used for qPCR are listed in table S1.

### Bioinformatic analysis of RNA-seq

RNA was sent to Novogene for RNA sequencing. Libraries were prepared using Illumina’s mRNA-Seq Poly(A) selection protocol and sequenced on a NovaSeq platform (paired-end 150 bp, unstranded) with a depth exceeding 20 million reads per sample. Initial quality filtering was performed by Novogene prior to delivery of clean reads. Raw reads were filtered according to the following criteria: removal of reads containing adapter sequences, removal of reads in which ambiguous nucleotides (N) constituted more than 10% of either read, and removal of reads in which more than 50% of bases had a Phred quality score ≤ 5.

The strandedness of the libraries was verified using *how_are_we_stranded_here* and read quality was assessed with *FastQC* (v0.12.1). Samples were aligned to the Genome Reference Consortium Mouse Build 39 (GRCm39) with Ensembl annotation release 104 using *STAR* (v2.7.9a). Gene quantification was computed using *HTSeq* (v.0.13.5) and normalization was performed using *DESeq2* (v.1.34.0) in *R* (v.4.2.2). Differentially expressed genes were identified using an adjusted q-value < 0.05 and absolute fold change > 1.5 (Benjamini–Hochberg correction). All heatmaps, principal component analyses and volcano plots were produced in *R* (v4.1.2) using *tidyverse* (v2.0.0), *dplyr* (v1.1.4), *tibble* (v3.2.1), *ComplexHeatmap* (v2.10.0), *ggplot2* (v4.0.0) and *ggrepel* (v0.9.3), then refined and exported using *circlize* (v0.4.16), *viridisLite* (v0.4.2) and *svglite* (v2.1.3).

Gene Ontology (GO) analyses were conducted using the **DAVID** functional-annotation platform (Database for Annotation, Visualization, and Integrated Discovery; https://david.ncifcrf.gov/tools.jsp), as described.

All analyses were executed within a reproducible environment managed through *Conda* (v4.8.3) to ensure version control and computational reproducibility and performed within *RStudio* (Build 467).

### Statistics

Statistical analysis was performed using GraphPad software (https://www.graphpad.com/), version 7.0, San Diego, CA, applying the t-Test when 2 groups were compared, and ANOVA analysis with a Tukey’s multiple comparison test when comparing >2 groups. For cavernoma volumes in figures 5 and 6, Welch’s ANOVA test followed by a Dunnett’s T3 multiple comparisons test was used, based on unequal variances of the different groups. A value of p<0.05 was considered significant. All graphs represent the mean ± SEM with dots indicating individual values.

### Study approval

All procedures were performed in accordance with the European Parliament and of the Council Directive 2010/63/EU and Spanish legislation (RD 53/2013) and were approved by the Ethics Committee on Animal Welfare of University of Santiago de Compostela (reference 15010/19/003).

### Data availability

All individual data values represented in graphs are available in the Supporting Data Values file. The RNA-seq data is available in the NCBI GEO repository (accession GSE298723). RNA-seq differential expression bioinformatic analysis and figure creation are available at https://github.com/dvidmd/GSE298723_Differential_Gene_Expression_Analysis_Garci a_Colomer_et_al

## Authors’ contributions

C.M.P and J.Z. conceived the project. M.G.-C., J.E.M., M.S., A.G.-D. and D.G. set up the cellular and animal model and performed the experiments. L.D.-G. analyzed brains by micro-CT and measured lesion volume. E.M.E.-R., C.R., D.M., M.F. and M.V.-R. performed bioinformatic analysis of RNA-seq. C.M.P. and J.Z. drafted the manuscript. All authors read, discussed and approved the manuscript.

## Acknowledgments

The authors thank Dr Ralf Adams for generously sharing the (*Cdh5*) CreERT2 allele and Elisabetta Dejana for generously sharing the *Pdcd10*^fl/fl^ allele and the protocol for mBMEC isolation and culture. We also thank the Galicia Supercomputing Centre (CESGA) for the availability of computational resources.

The graphical abstract and schemes in figures 5 and 6 were created with Biorender.com.

## Supplemental materials

Figures S1 and S2.

Tables S1, S2, S3.

