## Supplemental information for "Rapamycin reverts cavernoma endothelial cell phenotype and when combined with lapatinib ameliorates chronic lesions"

### Supplementary material

#### Rapamycin can revert CCM phenotype in brain endothelial cells and ameliorates chronic cavernoma development in combination with lapatinib

Mar García-Colomer, José E Martínez, Luis Díaz, Miriam Sartages, Eva M. Esquinas-Román, Cristina Riobello, David Martínez-Delgado, Diego González-Pérez, Aurora Gómez-Durán, Miguel Fidalgo, Marta Varela-Rey, Celia M Pombo, Juan Zalvide

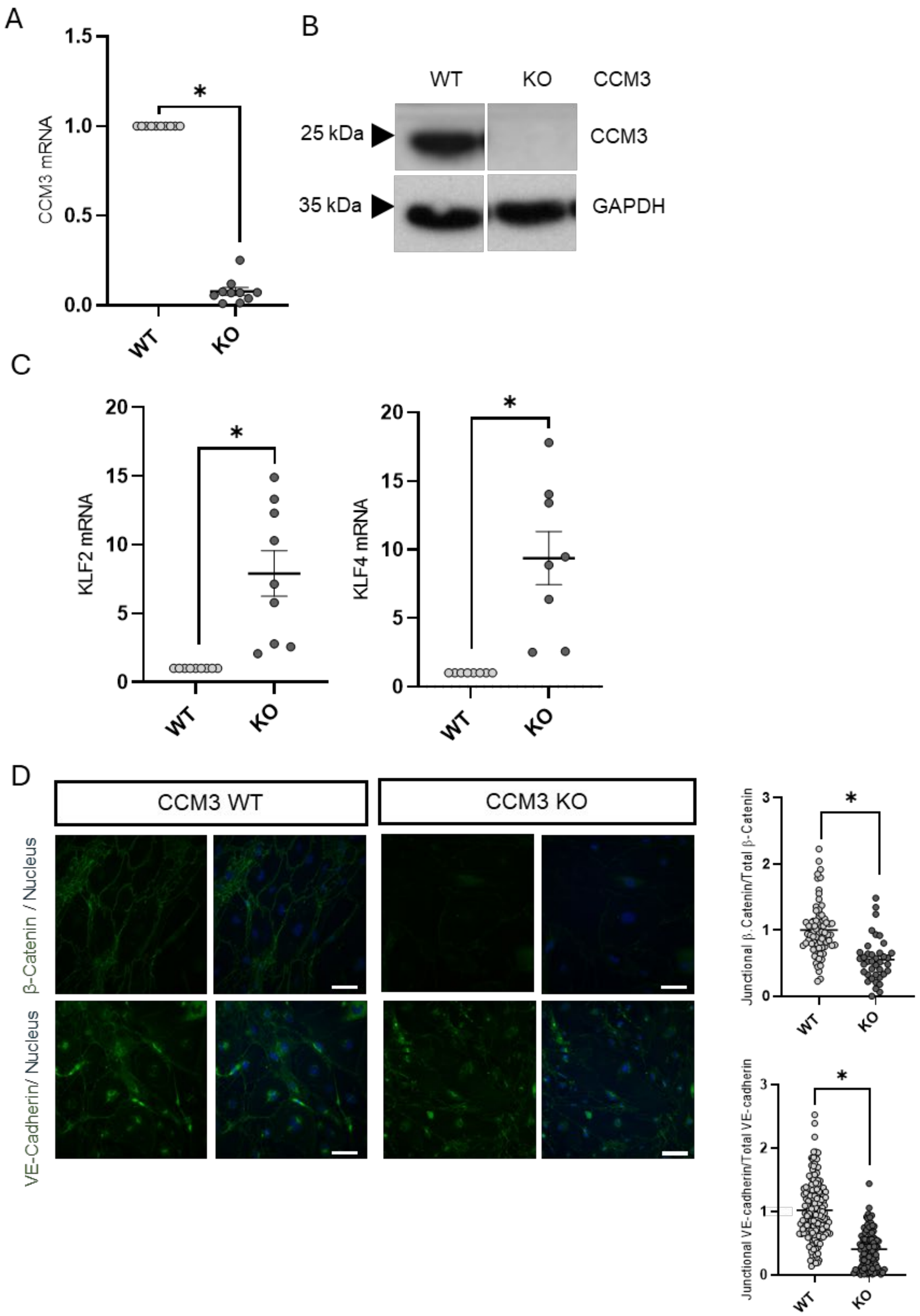
A
B
C
D
1.5
1.0
0.5
0.0
CCM3 mRNA
WT
KO
CCM3
25 kDa
35 kDa
GAPDH
KLF2 mRNA
KLF4 mRNA
20
15
10
5
0
CCM3 WT
CCM3 KO
β-Catenin / Nucleus
VE-Cadherin/ Nucleus
Junctional β. Catenin/Total β-Catenin
Junctional VE-cadherin/Total VE-cadherin
3
2
1
*

**Figure S1.** Loss of *Ccm3* in mBMEC induces *Klf2/4* upregulation and redistribution of β-catenin and VE-cadherin. mBMEC from PDCD10fl/fl cdh5(PAC)-CreERT2 mice were either treated with 4-hydroxytamoxifen to generate *Ccm3* KO cells (KO) or left untreated for *Ccm3* WT cells (WT). **A.** Downregulation of *Ccm3* mRNA in *Ccm3* KO mBMEC. An RT-qPCR was performed on *Ccm3* WT and KO cells of passage 2 and mRNA levels relative to GAPDH mRNA were normalized to those of *Ccm3* WT cells of each preparation. Graph shows the average and SEM of mRNA relative expression and the individual values of 10 independent biological replicates (cell cultures). Each cell culture was obtained from a pool of 3 to 4 brains. P-Value from Student's t test. **B**. Downregulation of *Ccm3* in *Ccm3* KO mBMEC. Cells were treated as in A and western blots were performed in *Ccm3* WT and KO cells. Shown is a representative western blot. **C**. Upregulation of *Klf2* and *Klf4* in *Ccm3* KO mBMEC. Cells were treated as in A, and an RT-qPCR for *Klf2* (left graph) and *Klf4* (right graph) was performed from *Ccm3* WT and KO cells. mRNA levels relative to *Gapdh* mRNA were normalized to those of *Ccm3* WT cells of each preparation. Graph shows the average and SEM of mRNA relative expression and the individual values of 9 independent biological replicates (cell cultures) for *Klf2* and 8 replicates for *Klf4*. Each cell culture was obtained from a pool of 3 to 4 brains. P-Value from Student's t test. **D**. Redistribution of VE-cadherin and β-catenin in *Ccm3* KO mBMEC. Cells were treated as in A and an immunofluorescence performed for β-catenin or VE-cadherin, counterstaining with DAPI. Left panel: representative photographs of the immunofluorescence with staining for VE-cadherin or β-catenin (green) and DNA (blue). Scale bar is 50 μm. Graphs on right show the ratio of junctional/total β-catenin (upper graph) or VE-cadherin (lower graph) after each treatment, referenced to *Ccm3* WT mBMEC of >50 cells per treatment from 2 independent biological replicates (cell cultures). Each cell culture was obtained from a pool of 3 to 4 brains. P-values are from ANOVA analysis with a Tukey's multiple comparison test.

A

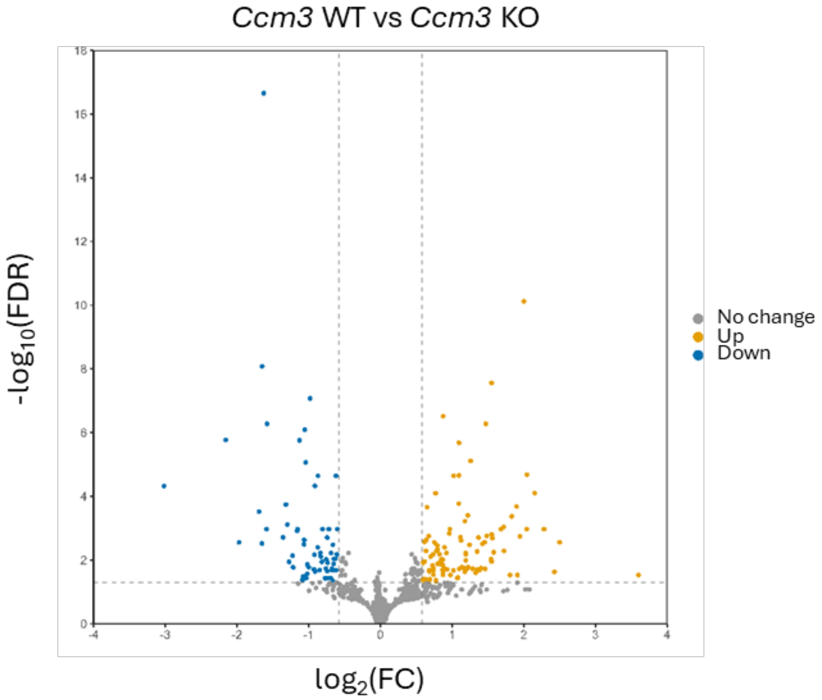


B

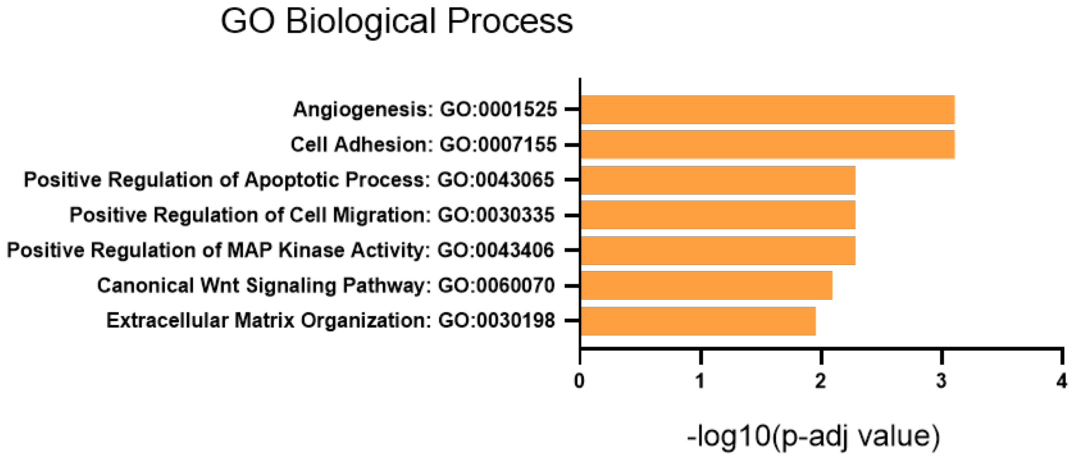


**Figure S2**: **A.** Volcano plot displaying the DEGs between CCM3 WT and CCM3 KO samples. Genes significantly upregulated or downregulated in KO compared to WT (KO/WT), with FDR < 0.05 and fold change (FC) > 1.5, are shown in yellow (Up) and blue (Down), respectively. Genes not meeting these thresholds are depicted in grey (No change). **B.** Gene ontology analysis of biological process of Ccm3 WT vs KO differentially expressed genes.

Table S1: Oligonucleotides used for qPCR

| Gene | Species | Forward/reverse | Sequence |
|---|---|---|---|
| *Gapdh* | Mouse | forward | AGGTCGGTGTGAACGGATTTG |
| *Gapdh* | Mouse | reverse | GGGGTCGTTGATGGCAACA |
| *β-Actin* | Mouse | forward | GTGACGTTGACATCCGTAAAGA |
| *β-Actin* | Mouse | reverse | GCCGGACTCATCGTACTCC |
| *Klf4* | Mouse | forward | GTGCCCCGACTAACCGTTG |
| *Klf4* | Mouse | reverse | GTCGTTGAACTCCTCGGTCT |
| *Klf2* | Mouse | forward | CTCAGCGAGCCTATCTTGCC |
| *Klf2* | Mouse | reverse | CACGTTGTTTAGGTCCTCATCC |
| *Nos3* | Mouse | forward | TGTGACCCTCACCGCTACAA |
| *Nos3* | Mouse | reverse | GCACAATCCAGGCCCAATC |
| *Ccm3* | Mouse | forward | TCACCGAGTCCCTCCTTCG |
| *Ccm3* | Mouse | reverse | GCCCGTGCCTTTTCATTTAGG |
| *Thbs1* | Mouse | forward | CCTGCCAGGGAAGCAACAA |
| *Thbs1* | Mouse | reverse | ACAGTCTATGTAGAGTTGAGCCC |
| *Adamts1* | Mouse | forward | AAGGAAGAAGCGATTTGTGTCC |
| *Adamts1* | Mouse | reverse | CCACCGAGAACAGGGTTAGA |
| *Dll4* | Mouse | forward | TTCCAGGCAACCTTCTCCGA |
| *Dll4* | Mouse | reverse | ACTGCCGCTATTCTTGTCCC |
| *Dll1* | Mouse | forward | GCAGGACCTTCTTTCGCGTAT |
| *Dll1* | Mouse | reverse | AAGGGGAATCGGATGGGGTT |
| *GAPDH* | Human | forward | GGAGCGAGATCCCTCCAAAAT |
| *GAPDH* | Human | reverse | GGCTGTTGTCATACTTCTCATGG |
| *β-Actin* | Human | forward | CATGTACGTTGCTATCCAGGC |
| *β-Actin* | Human | reverse | CTCCTTAATGTCACGCACGAT |
| *Klf4* | Human | forward | CCCACATGAAGCGACTTCCC |
| *Klf4* | Human | reverse | CAGGTCCAGGAGATCGTTGAA |
| *Klf2* | Human | forward | CTACACCAAGAGTTCGCATCTG |
| *Klf2* | Human | reverse | CCGTGTGCTTTCGGTAGTG |
| *Nos3* | Human | forward | TGATGGCGAAGCGAGTGAAG |
| *Nos3* | Human | reverse | ACTCATCCATACACAGGACCC |
| *Ccm3* | Human | forward | GCCCCTCTATGCAGTCATGTA |
| *Ccm3* | Human | reverse | AGCCTTGATGAAAGCGGCTC |

Table S2: Primary antibodies used for immunofluorescence or western blots

| Antigen | Supplier | Catalog number | Technique |
|---|---|---|---|
| VE-cadherin | Abcam | ab205336 | Immunofluorescence |
| b-catenin | BD Biosciences | 610153 | Immunofluorescence |
| GM130 | cell signaling technology | 70767 | Immunofluorescence |
| pS473-AKT | cell signaling technology | 9271 | Western blot |
| AKT | cell signaling technology | 2920 | Western blot |
| CCM3 | Proteintech | 66440 | Western blot |
| eNOS | BD Biosciences | 610297 | Western blot |
| pS1177-eNOS | BD Biosciences | 612393 | Western blot |
| GAPDH | Santa Cruz Biotechnology | sc-47724 | Western blot |
| KLF2 | Proteintech | 23384-1 | Western blot |
| pS371-S6K1 | cell signaling technology | 9208T | Western blot |
| S6K1 | cell signaling technology | 9202S | Western blot |
| Vinculin | Merck Life sciences | V4505 | Western blot |

Table S3: CCM3KO vs CCM3WT differentially expressed genes.

| GeneSymbol | baseMean | log2FC | pvalue | FDR |
|---|---|---|---|---|
| Ltb | 62.72608227 | -3.016152098 | 7.04644E-08 | 4.75366E-05 |
| Kirrel3 | 170.8127084 | -2.156462523 | 1.20697E-09 | 1.70991E-06 |
| Prdm8 | 26.88137404 | -1.970818315 | 1.16549E-05 | 0.002790147 |
| Abca9 | 113.8141038 | -1.691495982 | 5.94565E-07 | 0.000300829 |
| Ptx3 | 421.2757781 | -1.650356872 | 1.32178E-05 | 0.003020263 |
| Six1 | 268.1662715 | -1.648206667 | 1.75088E-12 | 8.26822E-09 |
| Id3 | 1595.730328 | -1.626452219 | 1.56666E-21 | 2.21949E-17 |
| Slco1a5 | 304.3607642 | -1.58798441 | 3.1764E-06 | 0.001071429 |
| Glipr2 | 854.0871797 | -1.581934401 | 2.82767E-10 | 5.29307E-07 |
| Bmp2 | 3187.518068 | -1.35627342 | 6.78512E-06 | 0.001948627 |
| Adora2a | 303.3684404 | -1.317176633 | 3.18763E-07 | 0.000180636 |
| Edn1 | 7918.723118 | -1.295852534 | 1.73443E-06 | 0.000767864 |
| Bdnf | 147.3574486 | -1.271510043 | 8.50014E-05 | 0.011413404 |
| Cfh | 118.0051309 | -1.223157861 | 4.4814E-05 | 0.007297466 |
| Egln3 | 1474.219699 | -1.217221811 | 0.000145627 | 0.016910625 |
| Ankrd1 | 2001.9677 | -1.161011641 | 3.71696E-06 | 0.001224607 |
| Pfkfb3 | 1929.919945 | -1.154423555 | 2.65467E-06 | 0.001071429 |
| Lamb1 | 7399.706169 | -1.128989411 | 1.35996E-09 | 1.7515E-06 |
| Cdkl1 | 182.0705683 | -1.089149443 | 0.000570673 | 0.042107945 |
| Gm17501 | 145.6547355 | -1.07903341 | 0.00041399 | 0.03276535 |
| Pdcd10 | 514.5099946 | -1.072275998 | 0.000509532 | 0.038601793 |
| Gna14 | 172.0070299 | -1.063983171 | 1.45407E-05 | 0.003218717 |
| Cpm | 151.2568284 | -1.062238539 | 9.11325E-06 | 0.002347407 |
| Pcdh7 | 809.8095393 | -1.055709275 | 5.1589E-10 | 8.12067E-07 |
| Ilvbl | 1355.74661 | -1.039771714 | 8.50463E-09 | 8.60608E-06 |
| Col27a1 | 124.7402879 | -1.031318169 | 0.000484101 | 0.03707165 |
| Prnd | 1517.553224 | -1.022270787 | 0.000322825 | 0.027386038 |
| Pim3 | 4036.250219 | -1.018921114 | 0.000110867 | 0.013540063 |
| Mmp25 | 1414.248069 | -0.998108717 | 0.000141946 | 0.016757957 |
| 2610008E11Rik | 2021.128806 | -0.979601399 | 2.9824E-11 | 8.45032E-08 |
| Mafb | 281.9450012 | -0.926211497 | 0.000189633 | 0.019627702 |
| Zic3 | 1168.382225 | -0.918188722 | 4.14184E-05 | 0.006822964 |
| Ccn1 | 17206.15254 | -0.912165393 | 6.64354E-08 | 4.70595E-05 |
| Adm | 12819.64874 | -0.910450566 | 0.000248873 | 0.022884298 |
| Dpysl3 | 6298.624777 | -0.874041328 | 1.93907E-05 | 0.003981282 |
| Thbs1 | 60206.47635 | -0.870589141 | 2.9369E-08 | 2.26995E-05 |
| Dapk2 | 1670.080386 | -0.845336525 | 0.000205355 | 0.020607788 |
| Vgf | 376.0753933 | -0.83775253 | 3.29642E-05 | 0.005993965 |
| Nox4 | 9553.510491 | -0.830959583 | 4.83782E-05 | 0.007788339 |
| Ppp1r3b | 243.9526751 | -0.814527566 | 8.53971E-05 | 0.011413404 |
| Fblim1 | 1546.850782 | -0.805634662 | 2.84383E-06 | 0.001071429 |
| Cdh2 | 1034.498502 | -0.769580025 | 0.000495268 | 0.037722897 |

| | | | | |
|---|---|---|---|---|
| 4930555A03Rik | 111.8279767 | -0.759677448 | 0.000163682 | 0.0182589 |
| Ccn2 | 32489.27788 | -0.7546988 | 6.45553E-05 | 0.009526608 |
| Usp53 | 581.1385659 | -0.742317926 | 0.000237321 | 0.021974714 |
| Sgk1 | 2823.180809 | -0.741637884 | 7.23627E-06 | 0.001971466 |
| Ppp1r37 | 7620.38269 | -0.737884683 | 0.000479095 | 0.036887725 |
| Tnc | 9913.685685 | -0.735650592 | 9.40414E-05 | 0.012111675 |
| Kctd11 | 1011.621316 | -0.718622367 | 3.1038E-06 | 0.001071429 |
| Fzd1 | 792.2314361 | -0.694319096 | 0.000473614 | 0.036664998 |
| Prrg3 | 509.9895604 | -0.686553031 | 3.37719E-05 | 0.00601319 |
| Lratd2 | 199.3392226 | -0.677066059 | 0.000215617 | 0.020846255 |
| Tspan6 | 4564.824631 | -0.675661299 | 0.000175952 | 0.018884171 |
| Colec12 | 1846.702362 | -0.670553392 | 0.000213756 | 0.020846255 |
| Dach1 | 765.3075853 | -0.661267787 | 1.52002E-05 | 0.003312933 |
| Wnt9a | 1206.375339 | -0.660543844 | 0.000663359 | 0.046394946 |
| Unc5b | 3060.07656 | -0.658977556 | 0.000215192 | 0.020846255 |
| Sp5 | 1165.471768 | -0.649750414 | 0.000106812 | 0.01315836 |
| Arhgef37 | 278.2314833 | -0.647364899 | 5.96688E-05 | 0.009089547 |
| Col15a1 | 1332.739717 | -0.625722198 | 5.93036E-05 | 0.009089547 |
| Csrp2 | 2230.715987 | -0.619758713 | 2.98974E-08 | 2.26995E-05 |
| Pawr | 430.3480685 | -0.616073434 | 0.00022072 | 0.020846255 |
| Frzb | 4416.045847 | -0.607659138 | 3.92773E-05 | 0.006595663 |
| Ctps | 5456.07055 | -0.599117701 | 2.99432E-06 | 0.001071429 |
| Fam83g | 186.7768076 | 0.594435642 | 0.000544998 | 0.040423996 |
| Lgals3 | 1121.020195 | 0.597210206 | 0.000586256 | 0.042811828 |
| Ncoa3 | 1399.829307 | 0.599535509 | 9.34735E-05 | 0.012111675 |
| Slc9a3r2 | 18386.96167 | 0.599778849 | 8.52925E-05 | 0.011413404 |
| Tpcn1 | 3077.85531 | 0.616502744 | 1.07379E-05 | 0.002668835 |
| Ephb4 | 3510.055097 | 0.618689557 | 8.01631E-05 | 0.011102098 |
| Hsph1 | 4516.268083 | 0.62619507 | 0.00053478 | 0.039874914 |
| Gbp9 | 841.0182992 | 0.635944294 | 2.60845E-05 | 0.00513249 |
| Slc30a1 | 1844.929027 | 0.642743939 | 8.92746E-06 | 0.002342136 |
| Atp8a1 | 886.0448932 | 0.652329587 | 4.19146E-07 | 0.000219927 |
| Sec14l1 | 8976.630997 | 0.674702064 | 5.81146E-06 | 0.001751724 |
| Baiap2l1 | 265.8114902 | 0.681687555 | 0.000635902 | 0.045044143 |
| C2cd2l | 1933.500357 | 0.684023421 | 0.000206558 | 0.020607788 |
| Cbfa2t3 | 1280.891023 | 0.68453809 | 0.000253068 | 0.02297204 |
| Nt5dc2 | 400.1402335 | 0.690264742 | 0.0005248 | 0.039547056 |
| Itprip | 785.426925 | 0.692719902 | 0.000220506 | 0.020846255 |
| Kcnq1 | 1296.398527 | 0.700785452 | 4.8946E-05 | 0.00779121 |
| Ms4a6d | 508.7907815 | 0.728714098 | 0.000124091 | 0.01489831 |
| Sema3g | 1472.637219 | 0.738739443 | 0.000153205 | 0.017489535 |
| Lama5 | 6526.196252 | 0.739665865 | 3.81138E-05 | 0.006584856 |
| Plpp3 | 1372.492044 | 0.740444802 | 0.000347192 | 0.028933358 |
| Zbtb46 | 499.7826492 | 0.741496461 | 0.000215452 | 0.020846255 |
| Palmd | 1555.167919 | 0.755514644 | 1.21818E-05 | 0.002829182 |

| | | | | |
|---|---|---|---|---|
| Pcdh12 | 1064.532673 | 0.770252982 | 1.29514E-07 | 7.97748E-05 |
| Abi3 | 490.8464994 | 0.774114277 | 0.00060498 | 0.04328662 |
| Rps6kl1 | 414.0865163 | 0.775158171 | 0.000670128 | 0.04653778 |
| Mylip | 371.9005214 | 0.783434035 | 2.66156E-05 | 0.005165253 |
| Prodh | 373.4091244 | 0.79431125 | 1.65639E-05 | 0.003502391 |
| Gsdmd | 901.7368814 | 0.797348972 | 0.000106538 | 0.01315836 |
| Arhgef15 | 2354.556159 | 0.800665625 | 2.27801E-05 | 0.004610363 |
| Zfp697 | 1023.276333 | 0.827832534 | 3.93904E-05 | 0.006595663 |
| Tsc22d3 | 684.7707486 | 0.833621317 | 0.000103703 | 0.013117513 |
| Dhcr24 | 1106.425077 | 0.843308784 | 7.91159E-05 | 0.01109737 |
| Rtl8c | 357.1292908 | 0.84928604 | 6.56261E-05 | 0.00958479 |
| Jag2 | 3543.88804 | 0.854849071 | 6.33805E-05 | 0.009451694 |
| Ccm2l | 2256.063516 | 0.868449977 | 0.000199599 | 0.020343349 |
| H2-T23 | 1374.304328 | 0.870083953 | 6.6716E-05 | 0.009644539 |
| Apold1 | 8841.068485 | 0.871409154 | 0.000118212 | 0.014313809 |
| Rtl8b | 1883.415243 | 0.871915811 | 0.000254578 | 0.02297204 |
| Adamts1 | 3885.691865 | 0.87424367 | 0.000351949 | 0.029158257 |
| Slfn5 | 6589.197313 | 0.876875239 | 1.28608E-10 | 3.03664E-07 |
| Dok4 | 1071.392801 | 0.885242641 | 0.000154316 | 0.017489535 |
| Als2cl | 1580.216495 | 0.894422901 | 1.90624E-05 | 0.003971435 |
| Efemp2 | 549.4915741 | 0.920288201 | 3.3956E-05 | 0.00601319 |
| Tmc8 | 156.7903011 | 0.928554915 | 0.000189807 | 0.019627702 |
| Pik3r6 | 419.4333626 | 0.962036833 | 4.60579E-06 | 0.00148296 |
| Synm | 1197.687747 | 0.96975601 | 2.77563E-06 | 0.001071429 |
| Hspa1b | 202.1837333 | 0.975811059 | 0.000366251 | 0.029587535 |
| Gatm | 189.3256984 | 1.019613175 | 0.000228887 | 0.021379061 |
| Aatk | 491.6699674 | 1.022532157 | 3.04433E-08 | 2.26995E-05 |
| Gm41442 | 1639.89894 | 1.074521586 | 0.000468791 | 0.036490992 |
| Alas1 | 9658.049538 | 1.084001519 | 0.000450234 | 0.035435885 |
| Scd2 | 6279.979819 | 1.093645781 | 2.87882E-07 | 0.000169934 |
| Mmp15 | 568.4963918 | 1.096356858 | 2.48955E-08 | 2.20434E-05 |
| Sh3tc2 | 730.908145 | 1.097737543 | 1.77075E-09 | 2.09051E-06 |
| Podxl | 28731.25072 | 1.101674865 | 0.00017107 | 0.018848943 |
| Fam167a | 880.7669695 | 1.118431837 | 6.87734E-06 | 0.001948627 |
| Fam20a | 414.3809546 | 1.126168844 | 9.62344E-06 | 0.00243456 |
| Adgre5 | 2838.583467 | 1.137111565 | 0.00014171 | 0.016757957 |
| Magix | 53.30449114 | 1.159084375 | 0.000149169 | 0.017181077 |
| Lsr | 809.2030824 | 1.177975922 | 0.000257681 | 0.02310482 |
| Sema7a | 3331.687172 | 1.179534006 | 1.31904E-06 | 0.000602802 |
| Efr3b | 281.2883099 | 1.190239706 | 3.2097E-05 | 0.005983128 |
| Ramp2 | 10575.18011 | 1.193161467 | 7.15628E-05 | 0.010240706 |
| Ecm1 | 6559.986209 | 1.194066623 | 3.9573E-05 | 0.006595663 |
| Rtl8a | 2573.965051 | 1.20201259 | 0.000183461 | 0.019270305 |
| Ckb | 3249.226407 | 1.220852687 | 8.12732E-07 | 0.000397034 |
| Klf4 | 3046.532958 | 1.225087622 | 0.000143881 | 0.01684592 |

| | | | | |
|---|---|---|---|---|
| Zswim1 | 1126.193427 | 1.244347142 | 1.55355E-05 | 0.003334719 |
| Ccn3 | 438.7636686 | 1.258714945 | 7.09943E-09 | 7.73674E-06 |
| Nkd2 | 282.1308608 | 1.285588114 | 0.000171632 | 0.018848943 |
| Afap1l2 | 270.4064255 | 1.330855451 | 0.000287118 | 0.024954612 |
| Irx3 | 321.4973199 | 1.348923296 | 0.000197453 | 0.020270428 |
| Endou | 137.0963968 | 1.350455557 | 2.8216E-05 | 0.005401847 |
| Gbp4 | 341.9624425 | 1.368870801 | 7.1893E-06 | 0.001971466 |
| Rd3 | 48.5164922 | 1.379874153 | 0.000219173 | 0.020846255 |
| Nr4a1 | 1078.577525 | 1.402095971 | 0.000159827 | 0.017970338 |
| Mcoln2 | 249.7837029 | 1.426035441 | 1.36086E-05 | 0.003060201 |
| Nkd1 | 76.91876467 | 1.453636873 | 1.16749E-05 | 0.002790147 |
| Ptprr | 388.4450224 | 1.462280306 | 0.000178035 | 0.018964053 |
| Jam2 | 260.4125113 | 1.469726648 | 2.98896E-10 | 5.29307E-07 |
| Ankrd33b | 536.4047846 | 1.484915629 | 5.68095E-06 | 0.001749611 |
| Syt7 | 72.79132867 | 1.545361478 | 6.32106E-05 | 0.009451694 |
| Kctd12 | 935.4859748 | 1.551037575 | 7.75372E-12 | 2.74617E-08 |
| Ip6k3 | 143.7245893 | 1.5517161 | 8.07169E-05 | 0.011102098 |
| Pltp | 31156.88426 | 1.552492819 | 5.01967E-06 | 0.001580303 |
| Ceacam1 | 999.6169729 | 1.562899094 | 7.67515E-06 | 0.002051582 |
| Dhrs3 | 72.00069253 | 1.581325784 | 3.0208E-05 | 0.005706085 |
| Nqo1 | 1745.515181 | 1.6812623 | 3.12317E-06 | 0.001071429 |
| Neurl2 | 1448.766416 | 1.719418411 | 2.15241E-06 | 0.000924036 |
| Nostrin | 66.06477977 | 1.723865671 | 2.59335E-05 | 0.00513249 |
| Slco2a1 | 86.30435762 | 1.809189316 | 0.000364782 | 0.029587535 |
| Anpep | 548.5413985 | 1.832484068 | 9.0059E-07 | 0.000425288 |
| Apod | 14346.60761 | 1.900408314 | 3.8415E-07 | 0.000209318 |
| Sox11 | 25.53210769 | 1.913245461 | 0.000369756 | 0.029595099 |
| Ednrb | 243.8240763 | 1.946902596 | 6.14175E-06 | 0.001812711 |
| Pcp4l1 | 294.5938665 | 2.002051686 | 1.07757E-14 | 7.63297E-11 |
| Rtp3 | 99.71238319 | 2.041942959 | 3.01077E-06 | 0.001071429 |
| Clic5 | 876.5589343 | 2.044745734 | 2.22378E-08 | 2.10029E-05 |
| Xdh | 171.8462856 | 2.15188893 | 1.22636E-07 | 7.89723E-05 |
| Car5a | 32.53607324 | 2.283159367 | 2.98112E-06 | 0.001071429 |
| Wnt9b | 26.6991604 | 2.428623316 | 0.000265565 | 0.023368112 |
| Spata25 | 137.4643994 | 2.501603867 | 1.18168E-05 | 0.002790147 |
| Srarp | 11.07924726 | 3.600018765 | 0.000364453 | 0.029587535 |
